# Whole genome sequencing of 358 brown planthoppers uncovers the landscape of their migration and dispersal worldwide

**DOI:** 10.1101/798876

**Authors:** Qing-ling Hu, Ji-Chong Zhuo, Yu-Xuan Ye, Dan-Ting Li, Yi-Han Lou, Xiao-Ya Zhang, Xuan Chen, Si-Liang Wang, Zhe-Chao Wang, Jia-Bao Lu, Norida Mazlan, Huy Chung Nguyen, San San OO, Thet Thet, Prem Nidhi Sharma, Jauharlina Jauharlina, S.M. Mizanur Rahman, Naved Ahmad Ansari, Ai-Dong Chen, Zeng-Rong Zhu, Kong Luen Heong, Jia-An Cheng, Shuai Zhan, Chuan-Xi Zhang

## Abstract

The brown planthopper (BPH), *Nilaparvata lugens*, is a serious migratory rice pest, which is distributed in the broad area of the tropical and temperate Asian-Pacific region. However, we know little about key aspects regarding its evolution such as how they diverged and dispersed worldwide. By resequencing and analyzing 358 BPH genomes from 92 populations across the world, we uncover the genetic relationships among their worldwide populations and the history of their global dispersal. We recovered five genetic groups representing the major population structures. Of these, Australian BPHs were shown large genetic divergence with Asian BPHs; two distinct groups have formed in South and Southeast/East Asia that show strong genetic admixture in the southwest border regions of China and west Thailand with Myanmar; two local populations in Bangladesh and Fujian province of China, respectively, unexpectedly separated with surrounding populations. We also find the genetic similarity and closely phylogenetic relationships between majority of East Asian BPHs and Indo-china peninsula BPHs, indicating that Southeast Asia mainland is the major insect sources and overwintering sites for East Asia. Our study provides important molecular evidence to address BPH evolution and other key aspects of its biology such as insecticides resistance and rice varieties virulence.

## Introduction

Rice (*Oryza sativa*) is amongst the staple food of most Asian people for more than 5,000 years, and now more than 90% of the world’s rice is planted and consumed in Asia-Pacific region, supporting half of the global population there. The brown planthopper (BPH), *Nilaparvata lugens* (Stål) (Hemiptera:Delphacidae), is a serious migratory rice (*Oryza sativa L.*) pest, which causes a conservative estimation of more than US$300 million per year world wide^1^. BPH is widely distributed across the region west from Pakistan to north-central Japan and south through the Malay Archipelago to New Guinea and the Solomon Islands, and north-eastern Australia^1^. Habitats of BPH extend through such a wide area, however, critical questions regarding its evolution, such as where BPH originated, how the species diverged, and how populations dispersed around the globe are still largely unknown.

BPH could not survive severe winters in temperate East Asian region north of 25°N^2^, because of the low temperature and no rice planting in winter, but the infestation of BPHs still occurs by new immigrants annually^3^, which migrate hundreds or even thousands kilometers far away by the favor of monsoon. In the past few decades, atmospheric trajectories were used to track insect sources and overwintering sites in East Asian countries, and the results supported an origin from the north part of Indo-China peninsula^4–7^. However, majority of BPH populations reside in the tropical areas of South Asia, Southeast Asia, and Oceania perennially, migration patterns in these regions remain unclear, even some migration movements has been detected in India^8,9^, south Vietnam^10^ and Philippines^11^. Mitochondrial markers have been proved to be difficult to distinguish regional populations in Asia^12,13^. BPH strains in different regions show phenotypic variation. Subtropical and temperate East Asian macropters were more adapted to migration with longer pre-oviposition periods and stronger starvation tolerance^14^, while tropical Asian strains (Southern Vietnam, Thailand, Malaysia, Philippines) were more adapted to reproduction^15^. BPH strains in South Asia, Southeast Asia and Oceania showed distinct virulence reactions to some rice varieties^16,17^, which suggests the exists of genetic difference among regional populations.

Here, we investigate the migration, dispersal and evolution of BPHs by performing whole-genome sequencing of 358 BPH individuals from 92 populations worldwide.

## Results

We sequenced the whole-genome of 358 BPH individuals collected from 92 sampling sites (Fig.1a) throughout South Asia (SA), Southeast Asia (SEA), East Asia (EA), and Australia, covering the most countries where BPH potentially exists. We also sequenced two individuals of *Nilaparvata muiri* as the outgroup. A total of 5.05Tb high-quality data were obtained after quality control, with an average coverage depth of 11X for each sample. By mapping to the improved BPH reference genome, we identified 1.95 million single nucleotide polymorphism (SNP) variations that were used for further population genomic analyses.

**Fig.1.**
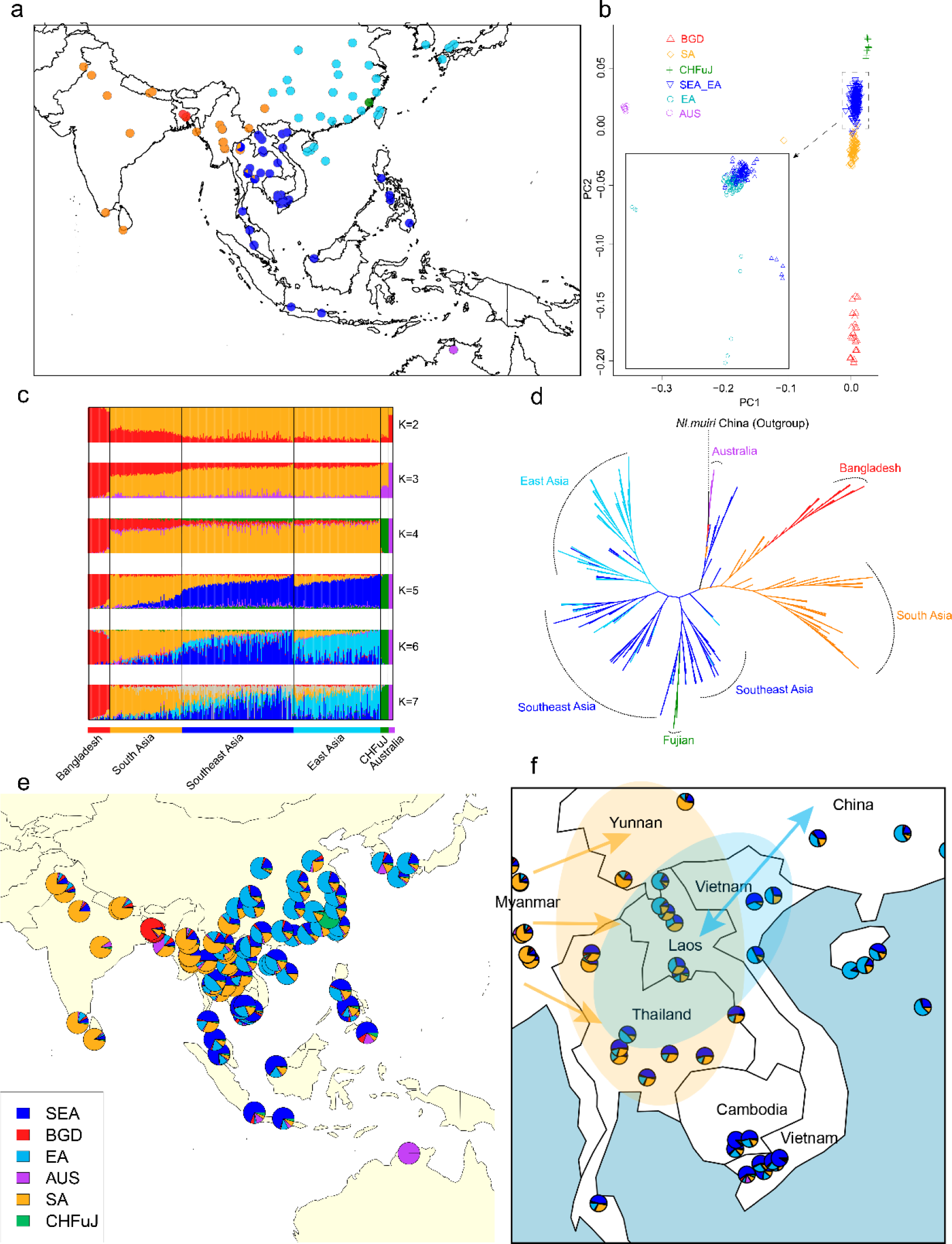
Population structure and phylogenetic relationships of BPH populations. **a** Geographic locations of sampling sites. **b** Principal component analysis (PCA) based on the first two components, inset shows separation of East Asia populations from Southeast Asia, BGD, Bangladesh; SA, South Asia; SEA, Southeast Asia; EA, East Asia; CHFuJ, Fujian,China; AUS, Australia. **c** ADMIXTURE analysis for K=2-7. Colored columns inferred the ancestry proportion of each sample. **d** Neighbor-joining tree based on 1000 bootstraps. **e** The average ancestry proportions for each sampling sites, according to ADMIXTURE analysis at K=6, which showing the genetic differentiation between EA and SEA. **f** The average ancestry proportions for each sampling sites, according to ADMIXTURE analysis at K=6, which showing the population genetic influence in SA and EA to SEA mainland.

### The genetic relationships among geographical populations

We analysed the genetic relationships among geographical populations based on the genome-wide SNPs. Principal-component analysis (PCA) unequivocally separated all BPH individuals into five groups (Fig.1b). Of them, there were two groups clustering multiple geographically-neighboring populations, i.e. the group SEA that, includes populations from seven Southeast Asian countries (Thailand, Cambodia, Laos, Vietnam, Maylaysia, Indonesia, Philippines) and three East Asian countries (China, Japan, South Korea) and the group SA that includes populations from four South Asian countries (India, Nepal, Pakistan, Sri Lanka) and a Southeast Asian country (Myanmar). We also found samples from Bangladesh, Fujian (China), and Australia form a distinctive group, respectively. Such an overall genetic grouping pattern was also well supported by the ancestry estimation and the phylogenetic analysis (Fig. 1c, d). AUS samples were shown relatively large genetic divergence with Asian populations, while the genetic divergence within Asian populations is low. The neighbor-joining (NJ) tree placed the FuJ clade as the sister clade to SEA and clustered the clades of BGD and SA together, agreeing with their geographic distributions. (Fig.1d). The well separation of these two populations suggests little gene flow of them with their surrounding populations which was probably caused by their unique local adaptions.

The NJ-tree further revealed that SEA group could be marginally subdivided into 3 subgroups, i.e. Southeast Asian mainland (SA mainland), Southeast Asian offshore (SEA offshore), and East Asia (EA) (Fig. 1d). Such a fine separation was also supported by the PCA analysis of SEA samples only (Fig. 1b). These results suggested gene flows frequently occurred among the populations of Southeast and East Asian region, and that the slightly genetic divergence between them was probably led by genomic loci that were under selection during annually migration in EA.

We found that some individuals in southwest Yunnan, the southwest border of China, west Thailand, and Laos showed close phylogenetic relationships with SA group individuals (Fig. 1d). Population structure analysis further revealed that populations in Yunnan, Thailand and Laos comprised large proportions of SA ancestry components (Fig. 1c,e). These observations suggested that regions in west Yunnan and Thailand might have received large amount of migrants from neighboring country Myanmar and the genetic structure of these populations are highly mixed.

All our population genetic analyses showed genetic relatedness between the majority of East Asian populations (except southwest Yunnan and Fujian) and SEA group (Fig. 1b,c,d). In the NJ-tree, except those individuals that formed EA subgroup alone, the rest of other East Asian individuals were undistinguishablely clustered with SEA mainland subgroup, indicating SEA mainland was the major yearly immigrant source of EA. In structure analysis at k=6, populations in northern Vietnam, north-central Laos, north-central Thailand showed more genetic similarity with East Asian populations (Fig. 1f). Thus, we hypothesized that these regions were the annually overwintering sites for East Asian populations.

Above results indicate a central role of SEA mainland in the genetic exchange of BPH, where migratory EA populations generally overwinter, populations on SEA islands frequently disperse, and migrants from west South Asian regions occasionally visit. SA mainland is an another place for genetic exchanges among regional BPH populations.

### Genetic diversity and demographic history

We found high levels of genetic diversities and individual heterozygosities in groups of SA, SEA (offshore and mainland), and EA, in consistent with the fast decay of linkage disequilibrium (LD) (Fig. 2a, b, c). These features suggest the high effective population sizes and frequently gene flows within these groups. In contrast, we found elevated LD and low level of genetic diversities and individual heterozygosites in three single populations, i.e. BGD, AUS and FuJ, indicating either founder effects existed in these groups or contracted population size (Fig. 2a,b,c,d). Tajima’D results also suggested that the BGD, AUS, FuJ groups have experienced population size contraction, whereas SA and SEA groups underwent expansion (Fig. 2b).

**Fig. 2.**
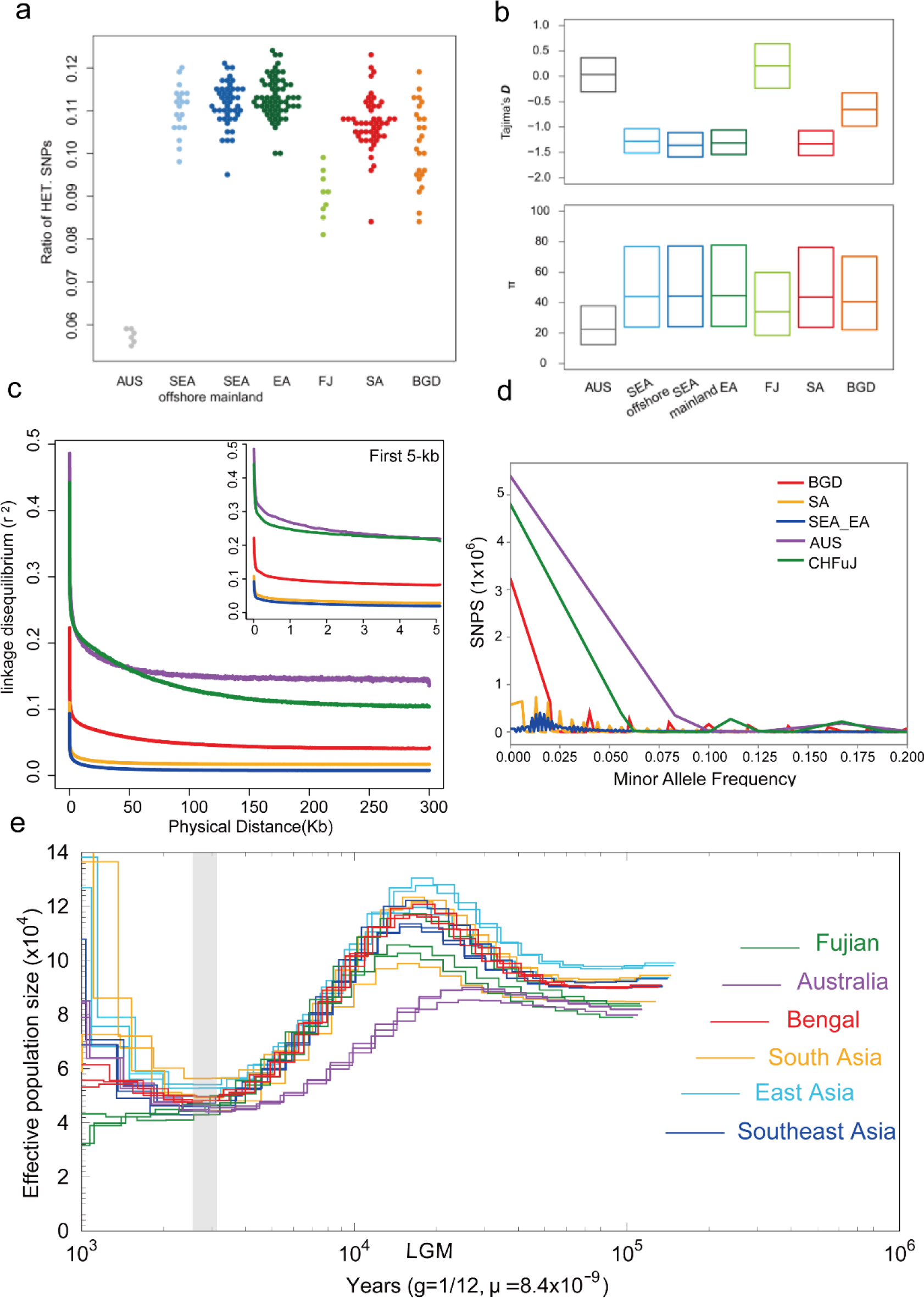
Genetic diversity and demographic history of BPH populations. **a** Heterozygosity of all 358 individuals, by calculating the ratio of heterozygous to homozygous SNPs. **b** Tajima’D and θπ scans for major groups using 10-kb windows on the genome. **c** Linkage disequilibrium (LD) decay for all the groups. **d** Minor allele frequencies (MAF) histogram. **e** Demographic history of the major populations inferred using PSMC. Each population was represented by 3 samples with the highest sequencing depth ranges from 12-fold to 17-fold.

We applied pairwise sequentially Markovian (PSMC) to estimate the historical size changes (Fig. 2e) and reconstructed a consistent demographic pattern across all main groups, except the slight deviation in the Australian population. BPH populations started to decline since approximately 20000 years ago until 2,000 years ago population sizes got recovered and reached the peak in about 1,000 years ago, in concordance with the earliest records of BPH outbreaks in 697 A.D in Japan^18^. These results supported a deep split between the Australian population and Asian populations, and that divergence between Asian populations occurred very recently. Previous study found the divergence of male courtship signals between Australian and Asian population, which influenced the success of hybridation, but showed little post-mating incompatibilities once mating successful^19^, reproductive isolation did not formed yet. The striking increase of population sizes reflected the intensified cultivation of domesticated rice in the past 2000 years.

## Discussion

Our studies clearly uncover the genetic structure, phylogenetic relationships, and evolutionary history among BPH geographic populations worldwide. Population structure and phylogenetic analyses revealed the existence of 5 distinct genetic groups and Southeastern Asia mainland was the major yearly immigrant source for East Asia. Our demographic analyses detected the populations expansion in recent 2,000 years in relation to the rice expanding cultivation.

The results revealed the deep divergence between Australian and Asian populations. Due to the low amount of insecticides input and host plant resistance selections, Australian populations might suffer a lower level of selection than Asian populations. However, we note that we only collected samples from wild rice in one site in Australia, which is probably not enough to represent the whole structure of Oceanian populations.

BPHs in Asia had formed two main genetic groups in South and Southeast Asia, which might be caused by the rice varieties, the different monsoon pattern between these two regions, the input of different types of insecticides, and the climate. Previous researches have reported the distinct virulence reactions to some rice varieties between South and Southeast Asian BPHs^17,18^. The genetic basis of the host virulence and local environment adaptation need further researches, and selective sweep scanning on the genome will be helpful.

Population structure analysis also showed massive genetic admixture in southwest Yunnan, west Thailand, and Laos. These populations will further migrate to East Asia and other part of Southeast Asia due to the massive migration within SEA group, all the samples in SEA and EA showed varying degrees of ancestry components from SA (Fig. 1c,e,f), which might suggested the gene flows between South and Southeast/East Asia were larger than we previous known^17^.

The data presented in this study will help us better understand the local adaptation of geographic populations that will be informative for the pest management.

## Methods

### Sampling

We sampled a total of 360 individuals, including 358 BPH individuals and 2 *Nilaparvata muiri* China. The East Asian population of BPH undergoes a yearly migration from the north Vietnam to Central-East China, Japan and South Korea. Our sampling sites for the migratory population covered the main stopover points from 16 provinces in 28 sampling points around China, and 3 points in Japan, 1 points in South Korea (Fig.1a, lightblue points). In southeast Asia, we sampled the tropical populations across seven countries, including 5 southeast Asian mainland countries, Vietnam (3 points in North region and 6 points in the South region), Laos (6 points ranging from north to south Laos), Thailand (9 points around Thailand), Cambodia (3 points in the South), Myanmar (7 points expand from Central to South regions) and 3 Southeast Asian offshore countries, Philippines (4 points from north to South), Indonesia (2 points), and Malaysia (4 points). In south Asia, our samples ranges in 5 countries, including Pakistan (1 point from the North East), India (4 points spanning North-Central to South), Sri Lanka(1 point), Nepal(2 points in Central) (Fig.1a, yellow points), and Bengal (3 points in Central and 1 point in Southeast Bengal) (Fig.1a, red points). We also sampled a site from north Australia to represent the BPH population in Australia (Fig.1a, purple point).

### Sequencing and quality control

Genomic DNA extraction, library construction and amplification followed standard protocols. All samples were sequenced on the Illumina sequencing platform (Hiseq X) with a pair-end read length of 150bp. We filtered raw data using the following thresholds to remove reads with low-quality and adapters: (1) unidentified nucleotides (i.e., N) more than 10% in a single read, remove the paired reads;(2) low quality bases (<=5) in a single read more than 50%, remove the paired reads;(3) remove reads aligned to adaptors. Finally approximately 33.78 billion clean reads were kept for later analysis.

### Mapping and SNP calling

Clean data were mapped against the BPH reference genome (updated 3^rd^ version) using Burrows-Wheeler Alignment Tool (BWA)^20^ v0.7.17-r1188 with command line ‘bwa mem ‒t 10 ‒k 32 ‒M ‒R’. The alignment results were further sorted by SAMtools v1.9. Consecutive variant calling were performed on GATK4 v4.1.3.0. Potential PCR duplicates were removed by ‘gatk MarkDuplicates’. Next, ‘HaplotypeCaller --emit-ref-confidence GVCF’ was applied to generate gvcf files for every individuals, and used ‘CombineGVCFs’ to combine the gvcf file of all the individuals and ‘GenotypeGVCFs’ was applied to call the raw variants sets. To reduce the SNP calling errors, we filtered out SNP variants according to the following thresholds: QD < 2.0, MQ < 40.0, FS > 60.0, SOR > 3.0, MQRankSum < −12.5, ReadPosRankSum < −8.0, QUAL<50, MAF < 0.05, --max-missing > 0.1, -max meanDP >30 --min-meanDP < 3. Potential variants were annotated by ANNOVAR software^21^.

### Population structure

Based on the genome-wide SNPs of 360 samples in our study, neighbor-joining tree were constructed using PHYLIP v3.697 with 1000 bootstraps. iTOL^22^ was used to present the constructed tree. PCA analysis was conducted by GCTA v1.92.1^23,24^. The population genetic structure was inferred using ADMIXTURE v1.3.0^25^ from k=2 to 7. We used TreeMix^26^ to infer the population split and mixtures.

### Genetic diversity and Demographic history

Linkage disequilibrium (LD) and minor allele frequency (MAF) among different populations were calculated by Samtools v1.9^27^. We estimated the heterozygosity for each sample by calculated the ratio of heterozygous to homozygous SNPs.

We inferred the historical population size of each population by pairwise sequentially Markovian coalescence (PSMC) model^28^. The generation time g was set as 1/12 year, and the mutation rate μ equal 8.4×10^−9^ as referred from Drosophila^29^.

## Acknowledgements

This work was supported by the National Natural Science Foundation of China (31630057 and 31871954). We thank Dr. M. Ashfaque (PARC, Pakistan), S. Zakaria(SKU, Indonesia), J. Pilianto (Brawijaya University, Indonesia), YH Song (GNU, Korea), Prof. H. Noda (NIAS, Japan), Dr. S. Sanada (NARO, Japan), Dr. N. Kado (Plant Protection Office, Shimane Prefecture, Japan), Dr. G. Bellis (Department of Agriculture, Fisheries and Forestry, Australia), Dr. S. Sarathbabu (Mizoran University Aizawl, India), Dr. W. Sriratanasak(Division of Rice Research and Development, Thailand); Ms. W. Janlapha (Prachinburi Rice Research Center, Thailand), LB Ma Rema & M Ibisate (Aklan State University, Philippines), Dr. SH Gu (Taiwan Museum of Natural Science), Dr. X-D Li(Yunnan Academy of Agricultural Sciences), D-W Zhang (Zunyi Normal University), Prof. B Chen(Chongqing Normal University), D Wang (Northwest A&F University), Prof. F-K Huang (Guangxi Academy of Agricultural Sciences), Y-K Li(Hainan University), Prof. Y-Z Li (Hunan Agricultural University), C-Y Niu (Huazhong Agricultural University), Prof. L-H Lu (Guangdong Academy of Agricultural Sciences), Profs.T-Y Wei and G Yang (Fujian A&F University), Mr. J-l Peng (Protection Station of Xiamen, Fujian), Mr. Z-L Shu(Zhenjiang Institute of Agricultural Science, Jiangsu), Dr. S-L Jin (Xinyang Normal University), Prof. H-J Xiao(Jiangxi Agricultural University), Dr. H Ma(Shandong Academy of Agricultural Sciences), Profs. KL Heong, Z-Y Liu, M-X Jiang (Zhejiang University), W-Q Zhang(Sun Yat-sen University), Prof. Z-X Lu and H-X Xu(Zhejiang Academy of Agricultural Sciences), Mr. X-Y Zhou and Prof. K-M Wu (Institute of Plant Protection, CAAS) for their kind helps in BPH collections.

